# A Multi-State Birth-Death model for Bayesian inference of lineage-specific birth and death rates

**DOI:** 10.1101/440982

**Authors:** Joëlle Barido-Sottani, Timothy G. Vaughan, Tanja Stadler

## Abstract

Heterogeneous populations can lead to important differences in birth and death rates across a phylogeny Taking this heterogeneity into account is thus critical to obtain accurate estimates of the underlying population dynamics. We present a new multi-state birth-death model (MSBD) that can estimate lineage-specific birth and death rates. For species phylogenies, this corresponds to estimating lineage-dependent speciation and extinction rates. Contrary to existing models, we do not require a prior hypothesis on a trait driving the rate differences and we allow the same rates to be present in different parts of the phylogeny. Using simulated datasets, we show that the MSBD model can reliably infer the presence of multiple evolutionary regimes, their positions in the tree, and the birth and death rates associated with each. We also present a re-analysis of two empirical datasets and compare the results obtained by MSBD and by the existing software BAMM. The MSBD model is implemented as a package in the Bayesian inference software BEAST2, which allows joint inference of the phylogeny and the model parameters.

**Significance statement:** Phylogenetic trees can inform about the underlying speciation and extinction processes within a species clade. Many different factors, for instance environmental changes or morphological changes, can lead to differences in macroevolutionary dynamics within a clade. We present here a new multi-state birth-death (MSBD) model that can detect these differences and estimate both the position of changes in the tree and the associated macroevolutionary parameters. The MSBD model does not require a prior hypothesis on which trait is driving the changes in dynamics and is thus applicable to a wide range of datasets. It is implemented as an extension to the existing framework BEAST2.

## 1 Introduction

Model-based phylogenetic and phylodynamic inferences are widely used to study both epidemics and macroevolution by using genetic sequences to reconstruct evolutionary processes. In many cases, the underlying population is structured, i.e it is composed of many different subpopulations which are subject to different evolutionary dynamics. Ignoring this structure and the resulting lineage-specific changes in evolutionary parameters can lead to biases in the inferred phylogeny and parameter estimates [1].

Multi-state birth-death models have been widely used to model population structure and analyze phylogenies built from individuals in a structured population [2, 3, 4, 5], both in epidemiological and macroevolutionary applications. These models contain a series of discrete states with state-specific birth and death rates, such that each state corresponds to a specific evolutionary regime. Based on a phylogeny where each tip is associated with a state, the state-dependent birth-and death rates are estimated. Birth events correspond to transmission events in epidemiology and speciation events in macroevolution, while death events correspond to becoming-non-infectious events in epidemiology and extinction events in macroevolution. A state might be for example a geographic location or the presence of a particular trait.

The Binary State Speciation and Extinction (BiSSE, [2]) and its extension to multiple states MuSSE, included in the package Diversitree [3], were the first efforts to infer state-specific birth and death rates from ultrametric phylogenies, i.e trees with all tips sampled at the same point in time, where each tip is assigned to a state. In [4], these approaches were extended to non-ultrametric trees. More recently, the Beast2 package BDMM [5] allowed the joint reconstruction of a phylogeny and quantification of the parameters of an underlying multi-state birth-death model. These approaches all have in common that the model is conditioned on a particular total number of states and the state at each tip in the phylogeny. This necessitates the formulation of a hypothesis as to which underlying feature drives the pattern of evolutionary rates. The BiSSE models in particular have been criticized for their approach being biased towards inferring trait-dependent rates regardless of the chosen trait [6]. Although this was addressed by the introduction of the HiSSE model [7] which uses a more appropriate null hypothesis, testing multiple different traits or combinations of traits would still require a different run of the inference for each. Thus there is a clear need for models which do not make such strong prior assumptions on the process driving the changes in evolutionary rates.

The method Bayesian Analysis of Macroevolutionary Mixtures (BAMM, [8]) addresses these issues and is able to infer the number of states, assign each lineage of the tree to a state and estimate the birth- and death rate parameters associated with each state. However, its results have been called into question, as [9] identified issues regarding the calculation of its likelihood function and a strong dependency on the prior when inferring the number of states, as well as inaccurate diversification rates estimates. Some of those criticisms were addressed by [10], which showed that the simulation used in [9] contained a large number of shifts which only affected small clades of the phylogeny, making them difficult to detect. [10] also pointed out that the sensitivity to the prior decreased sharply when using the default settings of BAMM rather than the setting used by [9]. However, issues regarding the calculation of the extinction probability in the likelihood function used by BAMM have to our knowledge not been addressed. Moreover, the process of moving between states is not explicitly modelled by BAMM, which may be a contributing factor to the prior sensitivity observed in some situations. Additionally, BAMM assumes that each state emerges only once along the tree. It thus implicitly links the changes in birth and death rates to lineage-specific innovations with no innovation occurring more than once, which may not adequately represent situations where the rates are driven by environmental or geographic conditions, for instance.

In this paper, we present a new Bayesian method for inferring lineage-specific birth and death rates jointly with a phylogeny, using a multi-state birth-death model. This method infers the number and position of evolutionary regimes as well as the state change rate, and requires strong assumption with respect to the features driving the variation in birth and death rates. We validate the implementation of this new method and evaluate its performance on simulated datasets. We then use it to re-analyze two empirical phylogenies and compare the results to those obtained by BAMM on those trees. Finally we discuss the limitations of the method and planned future work.

## 2 Results

We developed and implemented the MSBD model as a package within the BEAST2 framework. It takes genetic sequences or fixed phylogenetic trees as an input. The output is the inferred trees (in the case of sequences) and an assignment of lineage-specific birth and death rates to all lineages in the tree. Changes of these rates may happen anywhere along a branch. In what follows, we will first show evidence for the correctness of our implementation in a simulation study. Then, based on simulations, we investigate the accuracy of our tool when estimating the rates and change times. Last, we will present the results of an analysis of a lizard and a hummingbird phylogeny.

### 2.1 Validation: sampling from prior

To assess the correctness of the implementation of our model, we compare the distributions obtained by simulating under the model and by running the MSBD inference without data. The distributions are expected to match if the model is correctly implemented.

The results of the simulations without extinction are shown in Figure 2. The distributions obtained by forward simulation and by sampling from the prior match perfectly for all statistics, which provides strong evidence that the MCMC method is implemented correctly.

**Figure 1:**
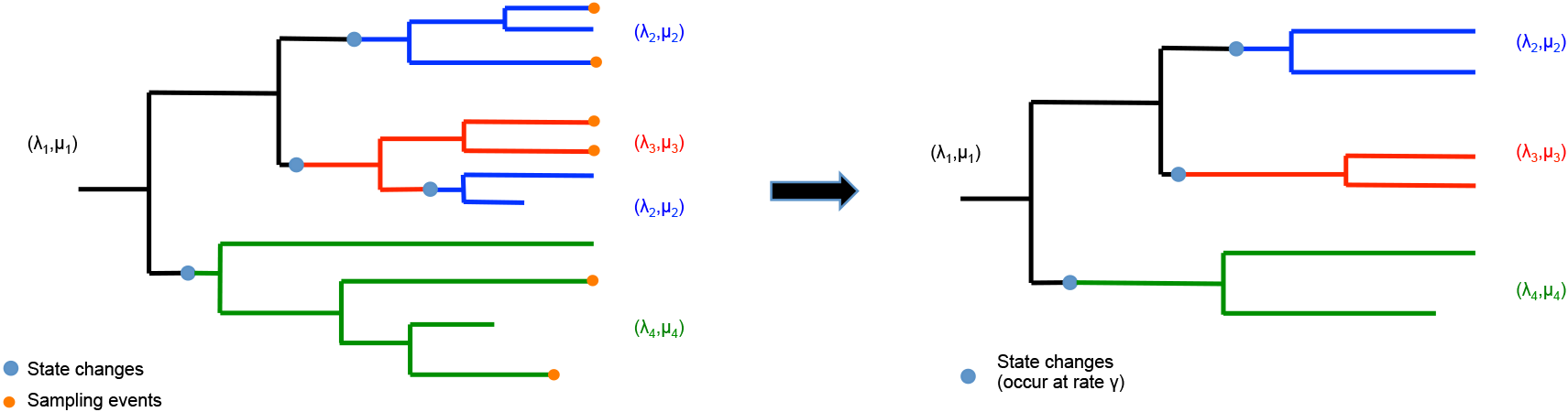
Visual representation of the multi-state birth-death model on a complete tree (left) with sampling events indicated in orange, and on the corresponding reconstructed tree (right). Each state is represented by a colour: the ancestral state, in black, starts at the root. The other states, in blue, red and green, start at change points along the tree. The same state can be present in multiple clades along the tree, such as the blue state in the complete tree.

**Figure 2:**
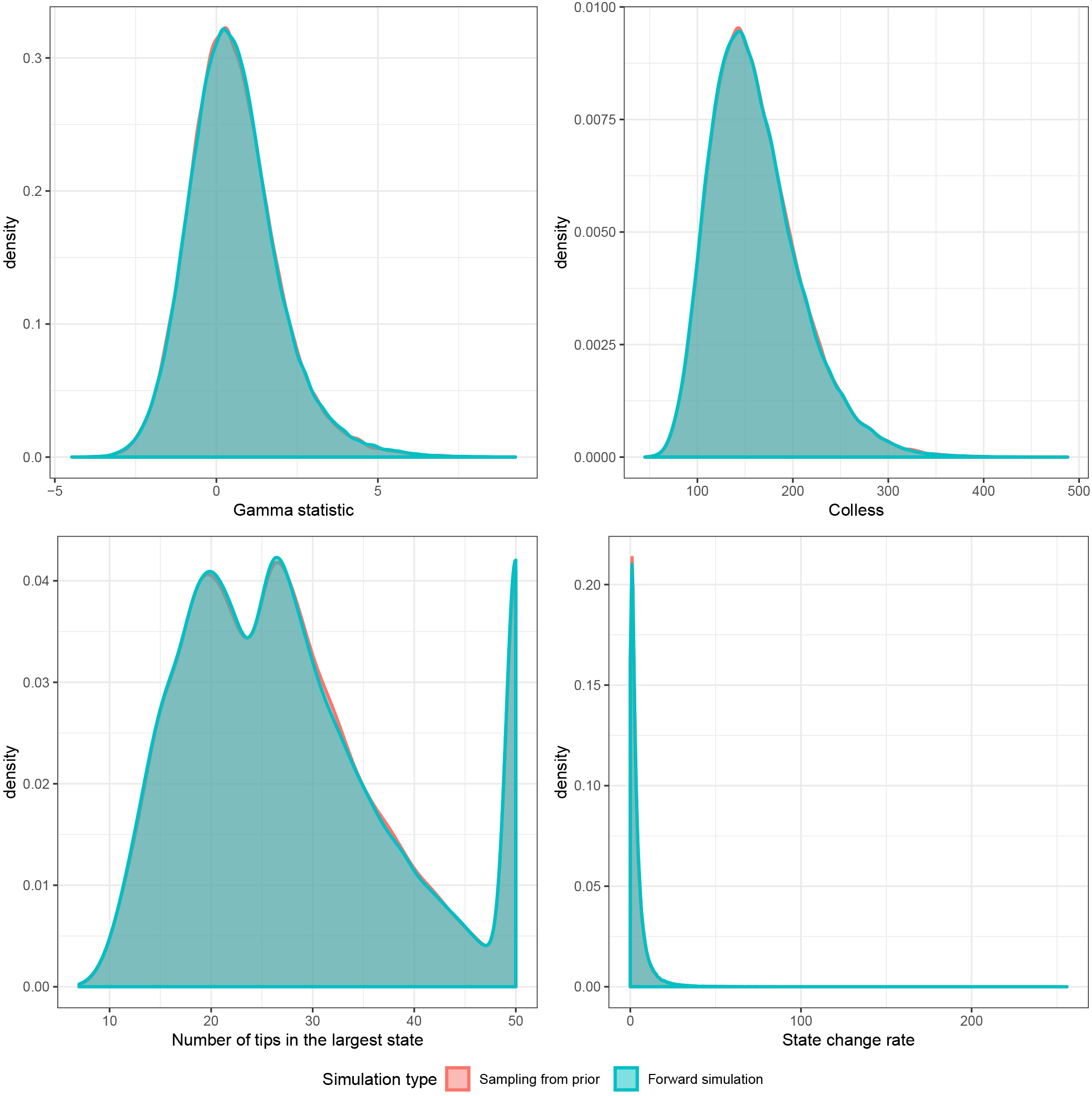
Comparison of the distributions of multiple summary statistics on trees obtained from forward simulation (in green) and MCMC sampling from the prior (in red) under a pure-birth MSBD process.

As expected, the simulations with extinction do not fully match between the two methods, as the forward simulation allows for state changes in the extinct parts of the tree whereas our method assumes there were none. As shown in Figure S1, there is a slight discrepancy in the statistics linked to the tree topology, and a stronger discrepancy in the statistics linked to the state distribution.

### 2.2 Accuracy of the inference

We use simulated phylogenies to assess the accuracy of the inference. Some datasets were simulated under the model, using a fixed set of states and a change rate γ. Other datasets were created by simulating two different trees under constant birth-death processes and attaching them. These joined datasets were thus characterized by the proportion *p* of tips in state 1 rather than γ.

#### 2.2.1 Parameter estimates

We evaluated the accuracy of the parameter estimates for the birth rates, death rates and state change rates by estimating the relative error of the median estimate and the coverage on simulated datasets. The error on the birth and death rates was evaluated both as an average across all tips and as an average across the entire tree, weighted by the edge lengths.

The results are shown in Figure 3. Estimates for the parameter γ are accurate for values around 0.2, corresponding to between 1 and 3 state changes in the tree on average. However, the estimates are much worse when γ is high, i.e. 2.61. This is likely due to the approximation of no state changes in the unsampled parts of the tree being more violated when γ is high.

**Figure 3:**
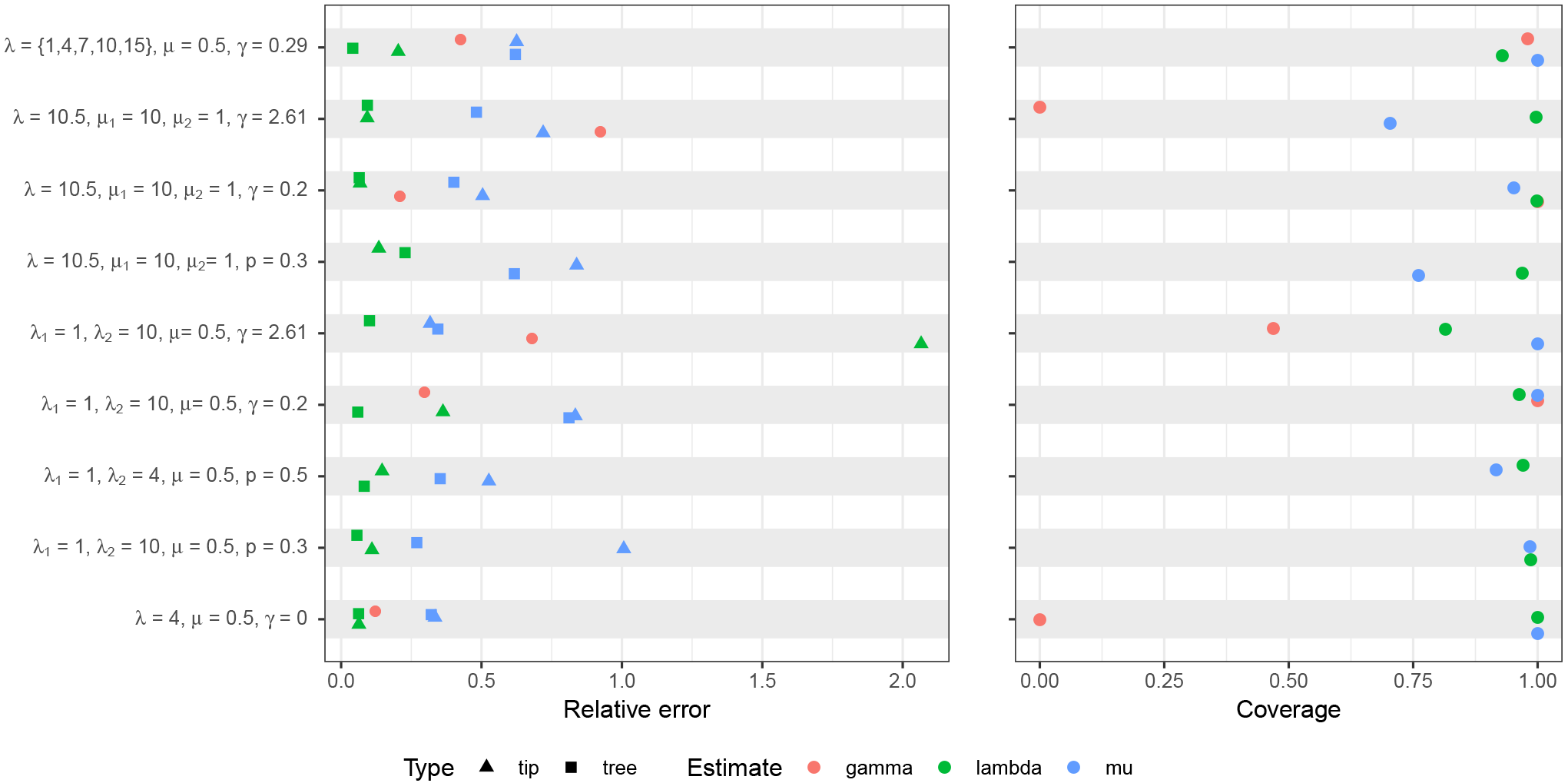
Performance of the birth, death and state change rates inference on different datasets. All measures are averages over 100 trees, with 200 tips for the datasets with 1 or 2 states and 500 tips for the dataset with 5 states.

Estimates of the birth rates are very accurate, except for the estimates at the tips under high γ. Since the estimates averaged over the whole tree do not suffer in a similar way, this exception is likely due to mis-attributing tips to the wrong regime rather than increased error on the regimes themselves: state changes affecting edges leading to tips, which are more likely when γ is high, cannot be detected by the inference, and will lead to tips being assigned a different state in the inference than the one recorded in the simulation. Estimates of the death rates are generally less accurate, although the true value is still in the 95% HPD interval in the vast majority of cases.

In conclusion, the MSBD method is able to recover the correct birth and death rates from simulated phylogeny, and is able to estimate the state change rate when it is small.

#### 2.2.2 State number and positions

We measure the accuracy of the inference regarding the number of states and the partition of tips into the different states. We use the Variation of Information (VI) criterion [11] to measure the distance between the inferred state placement and the truth: a measure of 0 indicates perfect concordance between the two. The upper bound of the VI distance depends on the number of states in the colouring and varies between 1.39 for 2 states and 3.22 for 5 states, however in this paper we have rescaled all VI distances so they range from 0 to 1, in order to make comparisons easier. VI distances were calculated for each sample of the posterior separately and on a “consensus” colouring built from the parameter values inferred for each edge. This consensus colouring put tips in the same state if the median estimate of their birth and death rates were less than 10% apart. Finally, we also estimated the posterior support for pairs of tips to be in the same state, split by whether these pairs are also in the same state in the true colouring.

Results are shown in Figures 4 and 5. The first finding is that the number of states inferred by MSBD is not a reliable estimate of the underlying process (Figure 4, right). In particular, the median estimate is similar for all datasets. Thus it should not be considered a good indicator of how many diversification regimes are in the process. The VI distance also shows discrepancies between the sampled clusterings and the truth on all datasets, in particular on the dataset with high γ, the dataset with 5 states and datasets with identical birth rate and different death rates (Figure 4, left). The consensus clustering however is closer to the truth on all datasets, which confirms that birth and death rate estimates are reliable. Due to the model, many simulated trees contains small clades of one state nested within another state. We expect these clades to be difficult to detect, as they cause small differences in the probability density. To test this hypothesis, we excluded all clades which contained less than 6 nodes (internal nodes included) from the true colouring, by attributing the tips of that clade to the ancestor state instead. Thus the tips belonging to small clades are not removed but simply recoloured (indicated as “With recolouring” in Figures 4 and 5). We observe a marked improvement in similarity when using this method, confirming that those small clades are unlikely to be detected by the MSBD inference. As seen earlier, the death rate estimates are less accurate than the birth rate estimates, and this is reflected by these results as well: the inference cannot easily distinguish between two states when when the death rates are different but the birth rates are identical even when those two states are clearly delimited in the tree (see row 4 of Figure 4). In conclusion, when states differ by their birth rates, the consensus colouring represents an accurate estimate of the original colouring, especially when excluding smaller clades. The quality of the inference is however much worse on states which only differ by their death rates.

**Figure 4:**
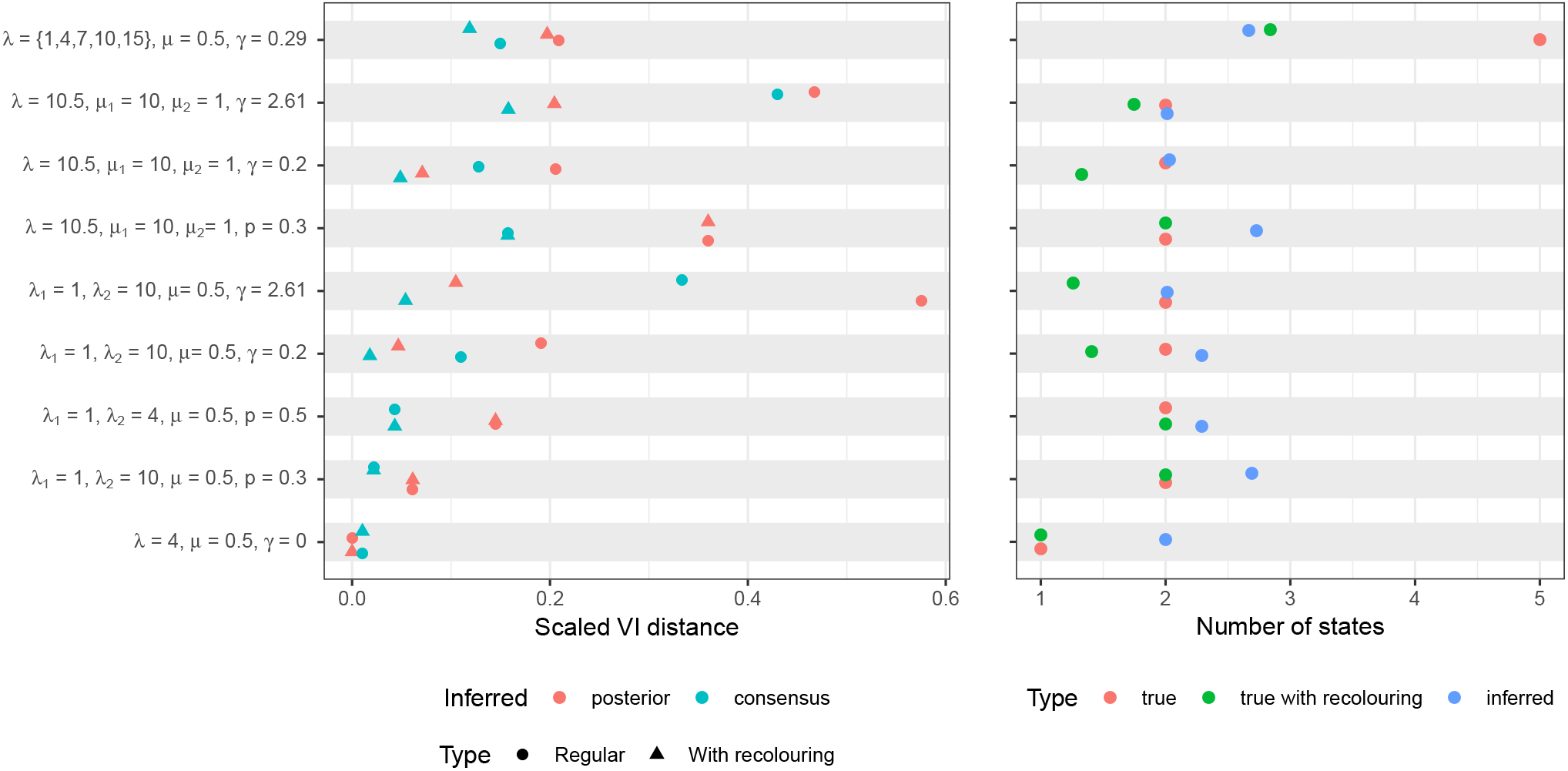
Performance of the state number and colouring inference on different datasets. All measures are averages over 100 trees, with 200 tips for the datasets with 1 or 2 states and 500 tips for the dataset with 5 states.

**Figure 5:**
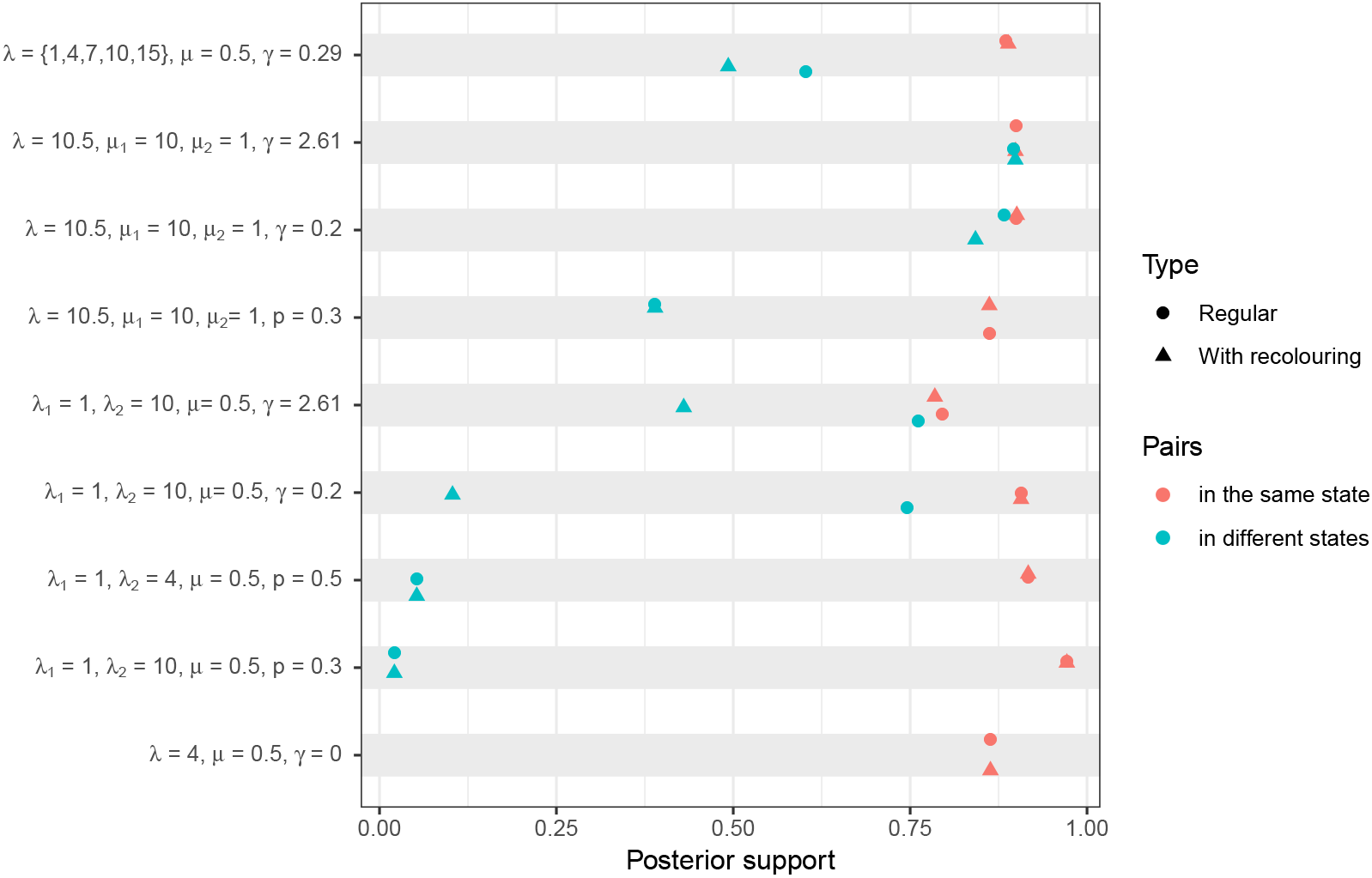
Posterior support for pairs of tips being inferred in the same state over different datasets. All measures are averages over 100 trees, with 200 tips for the datasets with 1 or 2 states and 500 tips for the dataset with 5 states.

We also looked at the posterior support for pairs of tips being in the same state, shown in Figure 5: if the inferred colouring is accurate, we expect pairs which are in the same state in the true colouring (in red in the figure) to have much higher support than pairs in different true states (in green in the figure). The results are consistent with the previous findings, showing that the posterior support reflects the true state partition if small clades are excluded and states have different birth rates.

#### 2.2.3 Tip state inference

Figure 6 shows an example of the posterior distribution on the birth rate for one tip of a tree. The tree was originally simulated with parameters λ_1_ = 1, λ_2_ = 10, *μ* = 0.5 and γ = 2.61. The figure shows a clear bimodal distribution, which is indicative that the inference has identified (at least) two separate diversification regimes across the tree, but that there is uncertainty on which regime this specific tip belongs to.

**Figure 6:**
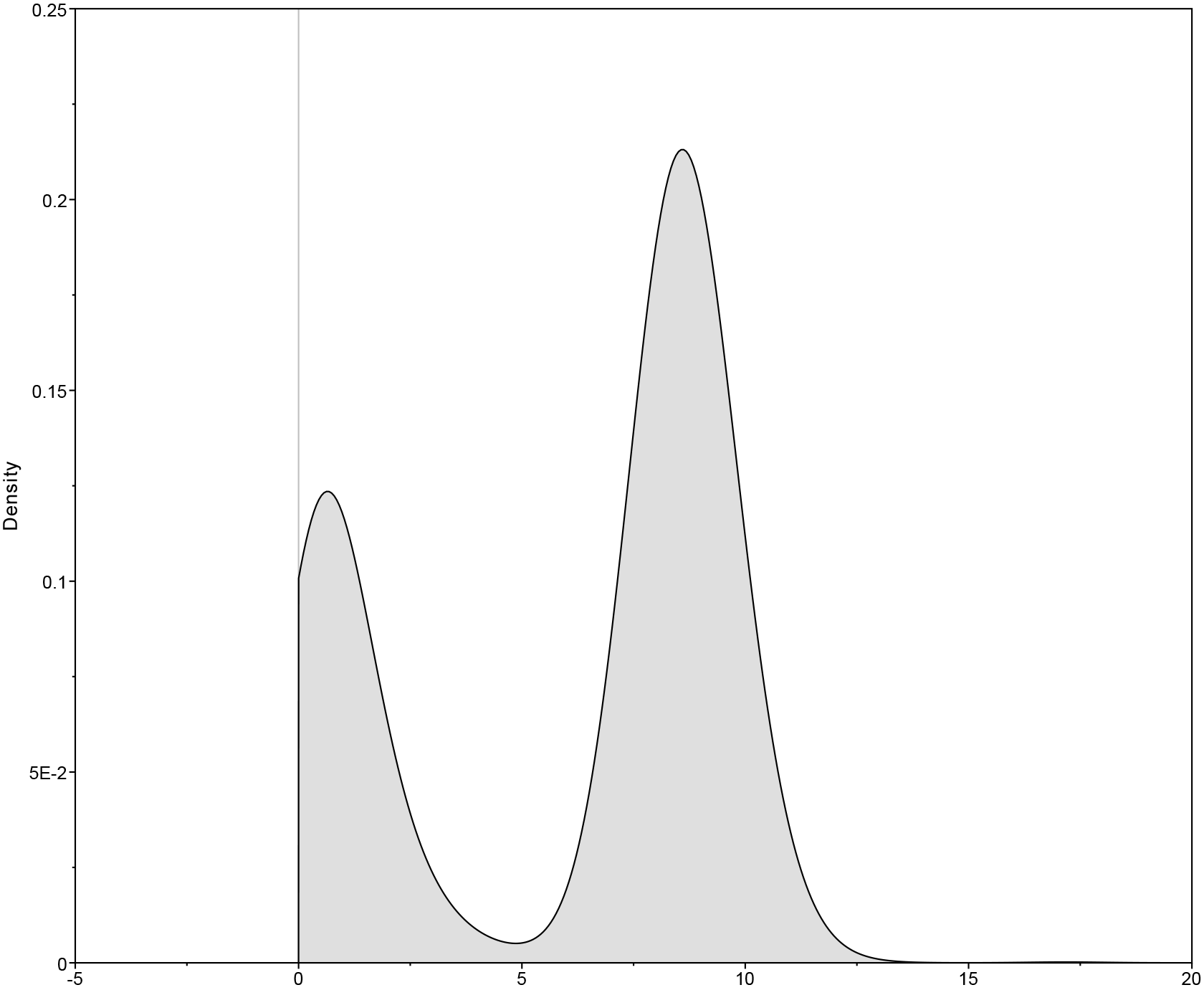
Bimodal posterior distribution of the birth rate on one tip of the tree, as inferred by the MSBD method.

This figure illustrates both the power of the MSBD inference, which is able to infer complex and nuanced evolutionary dynamics, and the complexity involved in interpreting the results. The median of the posterior is here 8.0, which corresponds to the most sampled state for this tip, but entirely misses the state with lower lambda. The 95% HPD interval is [0.0011; 9.94], which covers both states but gives no indication that the distribution is bimodal. Finally, the mean estimate is 6.0, which is a misleading summary of the distribution.

In this work we have used the median estimates to measure the accuracy of the inference, as it is the most representative of the configuration with the most posterior support. However, one should keep in mind that commonly used summary statistics can be flawed when summarizing distributions which are strongly multimodal.

### 2.3 Empirical datasets

We re-analyzed two empirical trees which were originally analyzed using BAMM: a phylogeny of hummingbird species obtained from [12] and a phylogeny of scincid lizards obtained from [13]. Both trees contain only extant species, with sampling proportions respectively *ρ* = 0.86 and *ρ* = 0.85. In both analyses, the sampling proportions were fixed to the truth and the priors for the birth and death rates were set to LogNormal(1.5,2.0). The tree topology was fixed and the prior on *n*\* was set to Poisson(4). The prior on γ was set to LogNormal(−4.0,1.0). We also performed a second analysis on the lizards phylogeny using the priors on birth rate and death rate which were originally used with BAMM, i.e Exponential(1.0) for both rates. Priors for γ and *n*\* were set to the same value as the previous analysis. The BAMM settings used on the hummingbirds phylogeny are not publicly available, so a similar analysis was not possible.

Average diversification rates per edge, weighted by the edge length, were logged for each edge. Figure 7, parts A and B, shows the results of the MSBD inference with lognormal priors on both empirical phylogenies, summarized as the median of the average diversification rate for each edge.

**Figure 7:**
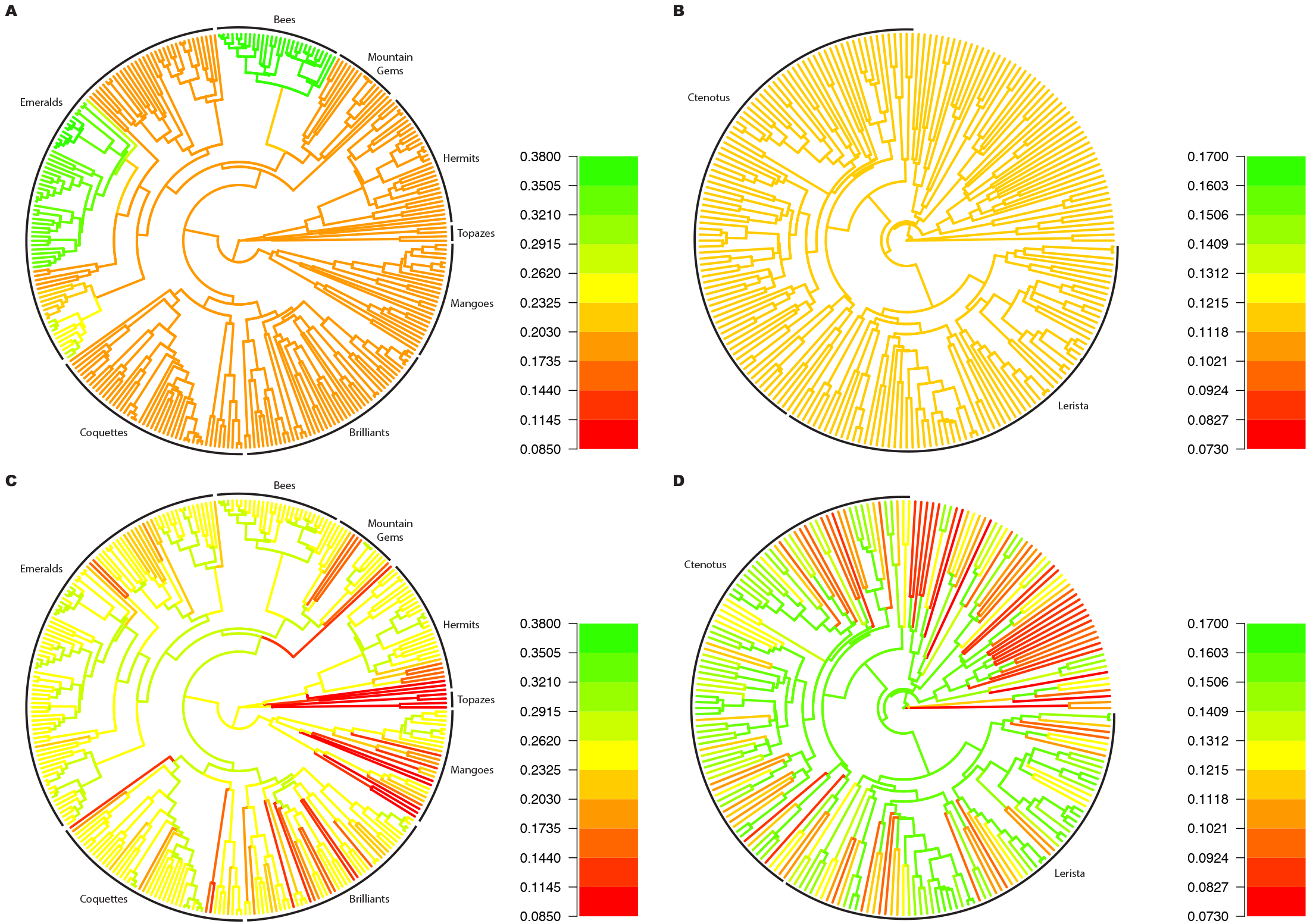
Empirical hummingbirds phylogeny (parts A, C) and lizards phylogeny (parts B, D) coloured by the median diversification rate inferred by MSBD for each edge. Inferences were run with a prior favoring low values of γ (parts A, B) or higher values of γ (parts C, D).

The hummingbirds inference (part A) shows some similarities with the original analysis by BAMM, but also differences. The diversification rates inferred by BAMM lied between 0.1 and 0.4, consistent with our results. BAMM also found strong posterior support for between 2 and 4 states with elevated diversification in the clade that includes Bees, Mountain Gems, and Emeralds, with particularly strong support for the Bees clade having a distinct diversification regime. In accordance with those results, the MSBD inference identifies 3 clades with elevated diversification rate, the Bees clade and 2 subclades of the Emeralds family. The main difference between the two inferences is that our method finds no evidence for time-dependency in the diversification rates, contrary to BAMM which infers an average speciation decay of 0.35 to 0.15 over 25 Myrs, corresponding to an exponential decay rate of 0.034 across the tree.

On the lizard phylogeny (part B), the results are quite different from the original analysis performed using BAMM. BAMM found strong support for two distinct configurations, one configuration with separate diversification regimes in the Lerista and Ctenotus clades and the rest of the tree, and one configuration with separate diversification regimes in the Lerista clade, the Ctetonus clade and the rest of the tree. MSBD on the other hand shows no evidence of separate diversification regimes in the tree, and infers a median speciation rate of 0.125 and a median extinction rate of 0.005 across the entire phylogeny. Similarly to the hummingbirds dataset, our method also detects no time-dependency in the diversification rate, although BAMM infers an average speciation decay rate of 0.2.

As time-dependency is not explicitly modelled in the MSBD inference, detecting it requires inferring widespread state changes across the tree. Thus the absence of time-dependency in our original inference could be due to the prior on γ being too low, and thus moving the inference away from this configuration. To test this hypothesis, we also ran an analysis with a much higher prior on γ, set to LogNormal(4.0,1.0). The results are shown in Figure 7, parts C and D. With the higher prior on γ, we can indeed recover signal for time-dependency in the lizards phylogeny, with edges close to the tips inferred to have a lower diversification rate than edges closer to the backbone of the tree. On the other hand, the hummingbirds phylogeny still shows no strong evidence for time-dependency, and no longer detects the clades identified as under different diversification regimes by the previous analysis. Thus it appears that when time-dependency is absent or weak, higher priors on γ can lead to a significant amount of noise and to the loss of signal for particular clades having different rates. This also illustrates the necessity of being careful when summarizing results from the MSBD inference, as a more in-depth analysis shows that edges in the hummingbirds phylogeny actually show a strong bimodal distribution which is very similar from edge to edge. The strong differences apparent in Figure 7, part C are in fact due to small variations in this bimodal distribution which lead the median to switch from one mode to the other.

Similar results were obtained when summarizing based on the median speciation and extinction rate, as well as when using the same priors as the original BAMM analysis. They are shown in Figures S2-S5.

## 3 Discussion

We have presented a new multi-state birth-death model for Bayesian inference of lineage-specific birth and death rates. The model is composed of multiple states, each associated with a specific birth and death rate, as well as a state change rate. The positions and times of state changes on the phylogeny then define to which state each lineage belongs to. The MSBD model thus represents a discretization of the true evolutionary process as a series of separate evolutionary regimes.

We have shown on simulated datasets that the MSBD inference can accurately estimate birth and death rates, and that those estimates can be used to build an accurate partition of the tree into states. However, our results also show that the MSBD inference cannot detect clades with different rates if the clades have very few tips. This is expected, as the method relies on the pattern of relative edge lengths to infer rates, thus small clades will not have enough signal to be inferred. Additionally, death rates estimates are less accurate than birth rates estimates in all simulation conditions. This in turn leads to lower accuracy in the inference of the colouring of the tree in datasets where states only differ by death rate, with many trees being inferred as presenting only one state.

The empirical analyses show two very different situations when using the default priors: on the hummingbirds dataset, our method and BAMM reach similar conclusions both regarding the presence and positions of separate diversification regimes and the parameter estimates. On the lizards phylogeny however, BAMM and MSBD obtain very different results, with MSBD finding no evidence of either rate changes or time-dependent rates. Further analyses on the empirical datasets show that MSBD is able to infer a pattern of time-dependent rates in a piecewise manner if there is signal for it, however this requires the prior for γ to be set much higher than for detecting single clades with different diversification regimes. It appears from our analysis that setting the prior in this way when no time-dependency is present can lead to noise and loss of signal. Thus one extension of MSBD will be to solve this issue by explicitly modelling time-dependent birth and death rates independently from changes in diversification regimes. It is to be noted that both empirical analysis were originally run with BAMM v1.0.0. BAMM has undergone significant changes since, including several bugfixes and modifications of the likelihood function, thus it is possible that the original results do not reflect the results which would be obtained with the latest version of the method.

One important thing to note is that interpreting the results of the MSBD inference requires more care than for other models, due to two primary reasons. The first is that the states are not linked to specific tips. If two MCMC samples contain *k* states, we cannot determine a precise correspondence between states in the first sample and states in the second. This problem is compounded by the variation in the number of states between different samples across the chain. Thus individual samples may not be a good representation of the overall inference. This is supported by the results shown in Figure 4, where the consensus clustering obtained from the rate estimates is much closer to the true clustering than the individual colourings sampled in the posterior.

The second reason is that the MSBD inference will frequently produce multi-modal posterior distributions on the rates associated with specific nodes or edges when the data shows signal for multiple regimes and there is uncertainty on which regime the node or edge in question belongs to. In these cases, the usual metrics used to describe Bayesian parameter estimates, i.e the median and HPD interval, give an incomplete picture of the output by failing to distinguish between uncertainty around the rate estimate and uncertainty on regime attribution. Thus analyzing the output of the MSBD inference should be tailored to the research question being considered, and may require different metrics than the ones we have used in this paper.

Future work will focus on explicitly implementing time-dependent birth and death rates to better accommodate situations where diversity-dependent or environment-dependent diversification is present, and on expanding the inference options available, in particular regarding sampling schemes. Currently only state-independent extinct and extant sampling are supported, and these are assumed to have known (fixed) values. We plan to incorporate a sampling scheme where each extant tip represents a genus or other group of species, as well as state-dependent sampling rates.

## 4 Materials and Methods

### 4.1 Multi-states birth-death model

We use a multi-states birth-death (MSBD) model with contemporaneous and non-contemporaneous sampling. This model contains *n*\* states, each associated with a specific birth rate λ_*i*_ and death rate *μ*_*i*_, *i* ∈ {1, 2,…,*n*\*}. The process starts with one individual in a state *r* picked uniformly at random from the *n*\* possible states, at time *t*_*or*_ > 0 in the past. Through time, each individual in state *i* undergoes birth events giving rise to an additional individual in state *i* with rate λ_*i*_, and dies with rate *μ*_*i*_. Additionally, each individual in any state *i* undergoes a change in birth and death rates to a different state *j* with rate *m*. Thus, the overall rate of change for any individual is γ = *m*(*n*\* − 1). Note that γ = 0 for *n*\* = 1.Throughout this paper, we consider γ (and not *m*) as a parameter.

The process stops at present time *t* = 0. The model includes both extinct and extant sampling: individuals are sampled upon death with a probability σ and individuals at the present are sampled with a probability *ρ*.

The process gives rise to complete trees, displaying all birth, death, state change, and sampling events (Figure 1, left). The reconstructed tree 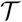 is obtained by pruning all lineages of the complete tree without sampled descendants (Figure 1, right). By analogy with the figure we will call the attribution of states to lineages and the position of state changes on the tree the colouring 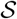 of the tree.

### 4.2 Probability density of a reconstructed tree

We derive the likelihood of the MSBD model on a given phylogeny, i.e the probability density of the reconstructed tree 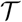 with the colouring 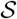, given the values of the birth and death rates for each state summarized in 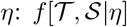.

We refer to a node in the phylogeny as either a branching event, a tip or a state change event. Thus the edges of 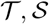, are the edges of 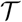 subdivided at state change events, and any edge belongs to only one state.

Following [5], we define *p*_*i*_(*t*) as the probability of a lineage in state *i* at time *t* > 0 not appearing in the reconstructed tree, i.e the probability of this lineage not being sampled before or at the present. We also define *q*_*i*,*N*_(*t*) as the probability density of a given edge *N* in state *i* at time *t* > 0 evolving according to the tree 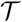 and states 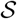 between time *t* and the present.

Note that 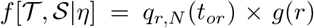, with *r* being the root state, and *g*(*r*) being the probability of the first individual being in state *r*. We assume here a uniform distribution, i.e. 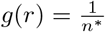.

In a similar fashion to [5], we obtain the ordinary differential equations Eq. 1 for *p*_*i*_(*t*) and Eq. 2 for *q*_*i*,*N*_(*t*) where *t* ∈ [*t*_*e*_;*t*_*s*_],*t*_*s*_ > *t*_*e*_ with *t*_*e*_ and *t*_*s*_ respectively the end and start times of edge *N*:

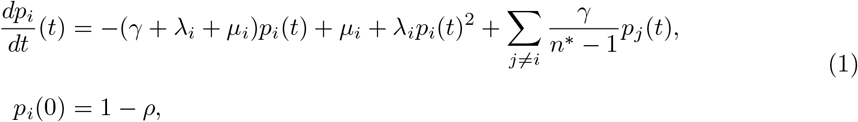

and

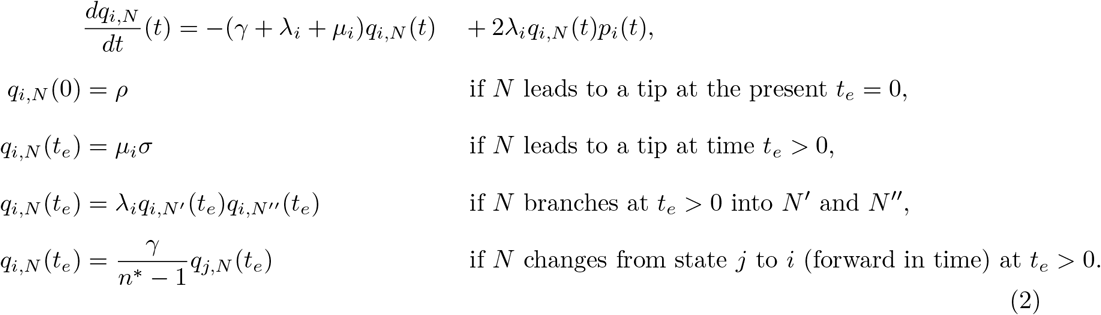

These ordinary differential equations do not have an analytical solution. Numerical integration is computationally expensive and can be unstable for certain parameters. Thus, in our implementation, we make the assumption that no state changes happen in the unsampled parts of the tree, meaning we observe all state changes in the reconstructed tree. With this assumption, the differential equation for *p*_*i*_(*t*) simplifies to Eq. 3.

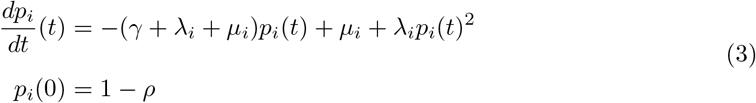

With this approximation we can derive an analytical solution for *p*_*i*_(*t*):

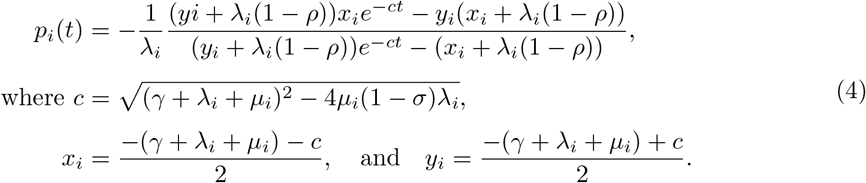

Using Equation (4) in the differential equation for *q*_*i*,*N*_(*t*) (Equation 2) allows us to derive *q*_*i*,*N*_(*t*) analytically:

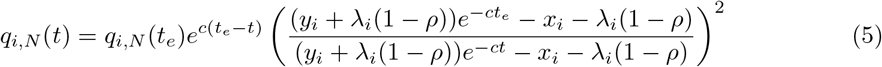

For an edge *N* in state *i* which starts at time *t*_*s*_ and ends at time *t*_*e*_ (*t*_*s*_ > *t*_*e*_), *q*_*i*,*N*_(*t*_*s*_) is the likelihood of the full subtree descending from edge *N*. The likelihood of edge *N* can be obtained as 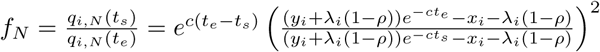.

This allows us to write the probability density of the phylogeny 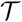 and the state changes assigned to the lineages 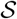, with *N*_*i*_ being the set of edges in state *i*, *B*_*i*_ being the set of birth events in state *i* (and for each event *b* ∈ *B*_*i*_, *t*_*b*_ the time of this event), *S*_*i*_ being the set of extinct tips in state *i*, *n*_*ext*_ being the number of extant tips, and *k* being the number of state change events:

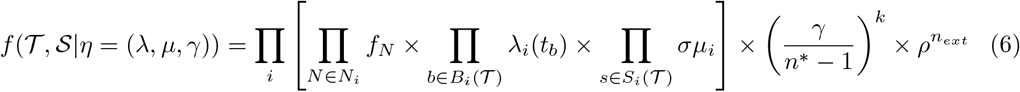

Note that if *n*\* = 1, then *k* = 0 and the term 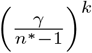 is removed from Equation 6. Note also that if the tree starts with 2 lineages at time *t*_1_ instead of 1 lineage at time *t*_*or*_, the likelihood becomes 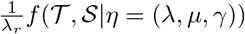.

### 4.3 Bayesian inference

We implemented our model in a Bayesian framework as an add-on to the popular MCMC inference software BEAST2 [14], which allows to estimate 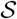 and *η* from a phylogeny based on Equation 6. The inference can be performed on a fixed tree 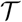, or directly on sequences, in which case 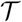 is inferred jointly with the other parameters using the substitution and clock models provided by BEAST2. In a joint inference, we sample from the following distribution:

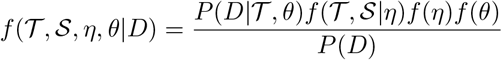

with the data *D* being the sequence alignment, *θ* being the parameters of the sequence evolution model, and *f*(*η*)*f*(*θ*) being the prior distributions for the model parameters, and 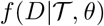, being Felsenstein’s likelihood for the sequencing data. If we condition on a fixed tree 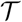, we use 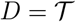.

While we infer *n*\* for our data, the number of states assigned to the reconstructed phylogeny, n, may be smaller than *n*\*, i.e. *n* ≤ *n*\*. To reduce the complexity of the computation, we do not sample the birth and death rates associated with the states which are not currently assigned to the tree, and instead marginalize over those rates. This marginalization introduces an additional term 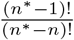 to the probability density to account for the sampling of *n*\* − *n* unassigned states.

It has been shown that in unstructured models, the three parameters λ, *μ* and *σ* are not identifiable [15]. In order to avoid potential parameter correlations in the structured model, we require the sampling probabilities *ρ* and *σ* to be provided as inputs.

More details on the implementation can be found in the Supplement.

### 4.4 Simulation study

To study the behaviour of our method, we simulated trees under our model using a range of parameter values. We used a stochastic forward in time simulation process which takes the following inputs:

- a stopping condition: the process is stopped upon reaching a certain number of tips or after a certain time had passed
- a rate γ of state change
- the total number of states in the process *n*\*
- a function to sample birth rates and death rates for all states
- sampling rates or sampling numbers for the extant tips and extinct tips

The birth-death process is started with either one or two lineages and is simulated with the Gillespie algorithm until the stopping condition is met or all lineages descending from one of the starting lineages have gone extinct, in which case the resulting tree is discarded. At the end of the process, lineages are discarded based on the sampling settings to obtain the reconstructed tree. If the sampling settings lead to no lineages being sampled, the resulting tree is also discarded.

#### 4.4.1 Validation: sampling from prior

To ensure that the implementation of our model is correct, we performed a comparison of the distribution of trees obtained from forward in time simulations of the process to the distribution obtained from running an MCMC inference without sequence data under our model with the same priors. This “sampling from the prior” procedure has been described in [16].

We performed two sets of simulations, one with death (i.e *μ*_*i*_ > 0 ∀*i*) and one without death (i.e *μ*_*i*_ = 0 ∀*i*). The distributions obtained without death should match if the model is correctly implemented, however we expect a discrepancy when simulating with death, due to the approximation made in the probability density function employed by the MCMC.

The number of tips was fixed to 50, and *t*_*mrca*_ was fixed to 1.0. The priors used were the following: LogNormal(1.5,1.0) for the birth rates λ_*i*_ and LogNormal(1.0,1.0) for the state change rate γ. The prior for the death rates *μ*_*i*_ was the Dirac function *δ*_0_ in simulations without death and LogNormal(−1.0,0.5) in simulations with death. The prior on *n*\* was set to Poisson(4).

The forward in time simulation was performed as follows. Parameters for five different states were drawn from the prior distributions, then a tree was simulated starting with two lineages in the same state, with this initial state being chosen uniformly at random. The simulation was stopped after a time *t* = 1.0, or when all lineages had gone extinct. The simulated tree was kept in the dataset if the following two conditions were met: the number of extant tips was *n* = 50 and the time of the most recent common ancestor *t*_*mrca*_ = 1.0, i.e neither of the original two lineages had gone fully extinct. New parameters were drawn from the priors for the next simulation, independent of the previous draw having resulted in a tree which was kept or not.

We assessed the match between the two distributions of trees on two measures: the gamma statistic, which measures the balance of recent branching events in a tree against older events, and the colless statistic, which measures the left-right balance of lineages in a tree. To assess the sampling of state positions we also compared the distribution for the number of tips in the state with the maximum number of tips and the number of sampled states.

#### 4.4.2 Accuracy of the inference

We assessed the quality of the MSBD inference on simulated datasets covering a range of possible configurations: constant birth and death rates, multiple states with different birth rates, multiple states with different death rates, and multiple states with different birth and death rates. Some of these datasets were simulated using the forward in time process described previously. Our parameter choices for λ and *μ* are displayed in Figure 3. In short, we performed one set of simulations with γ = 0. Then we performed a set of simulations with two different birth rates (λ_1_ = 1, λ_2_ = 10, *μ* = 0.5). Next we performed a set of simulations with two death rates, where the net diversification (birth-death) matched the simulation with the birth rate variation (λ = 10.5, *μ*_1_ = 10, *μ*_2_ = 1). The rationale for keeping the net diversification the same was to investigate the difference of performance of the method when varying birth vs. death rates in the light of as few changes as possible across the simulations. Finally we did a set of simulations with 5 birth rates and one death rate. We chose “low” and “high” values of γ such that the resulting trees would contain respectively between 1 and 3 state changes and between 10 and 14 state changes on average, excluding the changes on edges leading to tips. The “low” value was thus set to 0.2 for datasets with 2 states, and 0.29 for the dataset with 5 states, while the “high” value was set to 2.61.

This process often led to trees where one of the states only covered a small portion of the tree, and so there was little signal for the presence of two states. To address this issue, we also simulated so-called joined trees, which were made of two trees simulated separately under a constant birth-death process. The root of the smaller tree was then attached to the bigger tree such that the resulting tree was ultrametric. These joined datasets were thus characterized by the proportion *p* of tips in state 1 rather than by a change rate γ.

No sequences were simulated, and all analyses were performed with fixed tree topologies. Thus we estimated 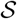 and *η* for a fixed tree 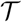. We measured the accuracy of the parameter estimates as well as the colouring 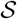.

## 6 Data accessibility

All simulations and analyses were done using custom R scripts. These scripts and the datasets are included in the Supplementary Materials. The method is publicly available as the BEAST2 package MSBD.

